# A novel 4D fMRI clustering technique to examine event-related spatiotemporal dynamics of face processing in naturalistic stimuli

**DOI:** 10.1101/2023.06.15.545143

**Authors:** C. Ekstrand

## Abstract

Cortical function is complex, nuanced, and involves information processing in a multimodal and dynamic world. However, previous functional magnetic resonance imaging (fMRI) research has generally characterized static activation differences between strictly controlled proxies of real-world stimuli that do not encapsulate the complexity of everyday multimodal experiences. Of primary importance to the field of neuroimaging is the development of techniques that distill complex spatiotemporal information into simple, behaviorally relevant representations of neural activation. Herein, we present a novel 4D spatiotemporal clustering method to examine dynamic neural activity associated with events (specifically the onset of human faces in audiovisual movies). Results from this study showed that 4D spatiotemporal clustering can extract clusters of fMRI activation over time that closely resemble the known spatiotemporal pattern of human face processing without the need to model a hemodynamic response function. Overall, this technique provides a new and exciting window into dynamic functional processing across both space and time using fMRI that has wide applications across the field of neuroscience.

---

Cortical function is complex, nuanced, and involves processing information in a multimodal and dynamic world. However, previous functional magnetic resonance imaging (fMRI) research has generally examined static activation differences between strictly controlled proxies of real-world stimuli that do not fully encapsulate the complexity of everyday multimodal experiences. For example, while it is well known that human face processing is a dynamic process, it has primarily been studied by comparing average brain activation between stimuli such as static Ekman’s faces (Dawel et al., 2021) and either scrambled faces or other object categories. Recently, in the growing field of naturalistic neuroscience, there has been a push to incorporate stimuli that better emulate conditions in the “real-world”. This field argues that to study “true-to-life” sensory experience, multisensory stimuli such as audiovisual movies, spoken or written narratives, and gaming environments, should be employed, as they better reflect the task demands encountered in everyday life. Indeed, it has been argued that capturing the complex, multiscale dynamics of the world around us is a primary organizational principle of the human brain (Gollo et al., 2015) and emerging evidence has suggested that naturalistic stimuli like audiovisual movies are preferentially processed compared to sparse stimuli. For example, Schultz and Pilz (2009) showed that, compared to static faces, dynamic faces led to more robust activation of face selective areas in the occipitotemporal cortex, including the putative fusiform face area (FFA). Thus, naturalistic stimuli provide exciting insight into how the human brain functions in more true-to-life conditions.

However, with this shift from strictly controlled experimental paradigms to more complex, true-to-life experiments comes the computational challenge of modeling, analyzing, and interpreting stimulus-response brain activation in a meaningful way. Several techniques have been proposed to analyze events in naturalistic data, each with their own strengths and weaknesses. A common approach to analyzing task-based fMRI data is reliant on the general linear model (GLM), which has also been applied to naturalistic data. For example, Häusler et al. (2022) investigated responses to spatial information in an audiovisual movie and an auditory narrative in the putative parahippocampal place area using GLM. Results from these analyses showed activation in the parahippocampal place area associated with spatial information in both types of naturalistic stimuli. However, these highly structured, event-based designs are not well suited for examining neural responses to complex, naturalistic stimuli as they impose specific time scale limits that may not represent the actual temporal and spatial response in everyday life (Sonkusare et al., 2019). Further, it is well-known that there are temporal correlations between adjacent time points due to sluggishness of the BOLD signal, machine drifts, physiological noise, etc., as well as spatial correlations between voxels, that make it difficult to accurately model the BOLD signal response (Ledberg et al., 2001). While temporal derivatives can be included to address small shifts in the HRF over time, the actual timing errors of the modeled HRF may be much larger (Bai et al., 2007). Of particular importance for naturalistic data, the canonical hemodynamic response function has been argued to not be reflective of the actual BOLD response to naturalistic stimuli (Polimeni & Lewis, 2021). For example, while the canonical response profile may be appropriate for brain regions involved in processing brief presentations of short and isolated events (e.g., areas near the primary auditory cortex), this does not necessarily extend to higher order brain regions (Ben-Yakov et al., 2012).

To alleviate problems with fitting the canonical hemodynamic response function to functional neuroimaging data, event-related averaging techniques that do not require explicit models of the hemodynamic response have been developed. A popular technique for modeling the hemodynamic response function related to specific events is the finite impulse response (FIR) technique, which estimates response amplitudes at each time point of a specific time window. FIR is more flexible than using a parametric filter shape (Goutte et al., 2000) as it is not biased towards the hemodynamic response function used in traditional fMRI analyses. However, one limitation of FIR models is the risk of overfitting the model due to the large number of parameters, which can lead to noise being modeled instead of signal. Further, it is difficult to model multiple stimulus conditions using FIR models (Bai et al., 2007), which is a limitation for the complex multisensory task demands of naturalistic stimuli. Indeed, Ben-Yakov et al. (2012) showed that while using impulse response models convolved with a standard hemodynamic response function to examine event-related responses to naturalistic sentences worked relatively well in low-level sensory regions (i.e., the primary auditory cortex), they did not accurately model context specific responses in higher level brain regions (such as the precuneus).

Ledberg et al., (2001) proposed a novel technique to analyze 4D fMRI data across experimental sessions that is of particular relevance to the current work. The researchers treated an entire 4D volume from each session as a single observation and modeled the time course over multiple sessions using a linear model. This generated statistical images that can be analysed using both parametric and non-parametric tests. They found that application of this method to fMRI data showed similar results to traditional 3D fMRI analysis methods. Of particular importance, the 4D clusters generated describe both spatial and temporal information about the BOLD signal response. This technique has the advantage of not relying on an explicit model for the BOLD contrast. However, one limitation is the need for multiple task-based sessions to fit the linear model, making it difficult to apply to naturalistic data where participants are required to watch long movies in the scanner. Further, repeated presentation of the same stimulus may alter the task demands with each subsequent presentation.

## The Current Study

Herein, we report a novel method for examining spatiotemporal dynamics associated with events in naturalistic stimuli that uses open-source, flexible, and widely used analytic tools by taking advantage of clustering across both space and time. To do so, we restructure fMRI data into 4D volumes associated with specific events in the audiovisual movies, in this case 10-seconds after either the onset or offset of human faces, and use each of these 4D volumes as a single observation. This is in contrast to traditional techniques that characterize data as 3D volumes obtained at a number of points in time. We then average multiple face-onset and face-offset 4D volumes to provide an estimate of the response amplitude for each time point.

Following this, we use spatiotemporal clustering to identified clusters of activation associated with face-onsets versus face-offsets in an audiovisual movie. Of primary importance, this technique does not require an explicit model of the BOLD signal response and takes advantage of both spatial and temporal autocorrelations to identify clusters of activation. Further, this highly flexible method can be applied to both asynchronous and synchronous naturalistic data without the need to specify GLM model parameters.

In this study, we use human face processing in naturalistic audiovisual movie stimuli as a test case for examining spatiotemporal dynamics, as face processing has been well characterized (see Duchaine & Yovel, 2015; Freiwald et al., 2016). Briefly, human face processing has been shown to recruit several core areas: the FFA, occipital face area (OFA), posterior and anterior superior temporal sulcus (pSTS and aSTS, respectively), ventral anterior temporal lobe (vATL), and the inferior frontal gyrus (Collins & Olson, 2014; Duchaine & Yovel, 2015). The FFA is located in the lateral middle fusiform gyrus and is thought to be primarily involved in holistic processing of faces. The OFA is located on the inferior surface of the occipital gyrus and is suggested to process low-level perceptual information of faces. The pSTS and aSTS are involved in processing changeable aspects of faces, including emotion, eye gaze, and lip movements. The vATL is thought to process higher-level, viewpoint invariant information (Collins & Olson, 2014). Finally, the inferior frontal gyrus is thought to mediate gaze perception and emotion. Facial motion (such as in audiovisual movies) elicits higher responses in face-sensitive regions such as the bilateral fusiform gyrus and left inferior occipital gyrus, as well as motion sensitive areas such as the posterior and middle superior temporal sulcus and area hMT+/V5 (Pitcher & Ungerleider, 2021; Schultz & Pilz, 2009). Regarding flow of activation, previous research suggests there is hierarchical processing from occipital to temporal areas. Using both fMRI and magnetoencephalography, Fan et al. (2020) found early activation of the right OFA and right posterior FFA, followed by activation in the anterior FFA and pSTS. Thus, based on previous research, we hypothesize that we will see early activation of the occipital and fusiform cortices, followed by flow of activation into the STS, that is greater for face onset than for face offset.

## Methods

### Data and Participants

This study used preprocessed data from 86 participants from the Naturalistic Neuroimaging Database (version 2.0.0). This database consists of structural MRI and fMRI data from 86 participants (42 females, ages 18-58, M = 26.81, SD = 10.09 years) who each viewed one of 10 feature length films during fMRI. All participants were right-handed, native English speakers, with no history of neurological/psychiatric illnesses or hearing impairments, unimpaired or corrected vision, and taking no medication (see Aliko et al., 2021 for more information). Full methods and preprocessing for these data can be found in Aliko et al. (2021). Briefly, fMRI data was acquired on a 1.5 T Siemens MAGNETOM Avanto MRI with a 32 channel headcoil using multiband EPI (TR = 1s, TE = 54.8 ms, flip angle = 75°, 40 interleaved slices, resolution 3.2 mm isotropic, multiband factor 4). The number of volumes ranged from 5470 to 8882 depending on the movie shown to the participants. Preprocessing was implemented with AFNI’s afni_proc.py pipeline (Cox, 1996; Taylor et al., 2018). For our analysis, we used preprocessed files that were not blurred or censored, with independent component analysis (ICA) based artifact time courses removed. Also included in the database was information about the timing of face onset and duration for each movie stimulus, as identified using machine learning via the AWS ‘Amazon Rekognition’ application programming interface (https://aws.amazon.com/rekognition/; see Aliko et al., 2021 for full details).

### Face Annotation Restructuring

The procedure for identifying Face-onset and Face-offset instances was as follows: first, we used the face annotation file provided by Aliko et al. (2021) for each movie stimulus to identify instances when a face was presented in the movie for at least 10 seconds. Second, to have the same number of Face-onset instances as Face-offset instances, we calculated the number of Face-offset instances in each movie that were at least 10 seconds long. Third, to choose Face-onset instances for each participant, we randomly shuffled the Face-onset array and selected the same number of Face-onset instances as total Face-offset instances. Face-onset instances were randomly chosen for each participant from all possible face onset instances in each movie. It is important to note that these instances were not controlled for face orientation, location, motion, emotion, distance to camera, number of faces, etc. and thus represent face processing across a wide array of audiovisual configurations. Face-offset instances were the same for each participant and each film, as face-offset from the screen for 10 or more seconds occurred less frequently than face-onset. Onset time and duration of each Face-onset and Face-offset instance were then saved as text files to be used in our analysis. The total number of instances for the Face-Onset and Offset conditions for each movie is shown in Table 1.

**Table 1.**
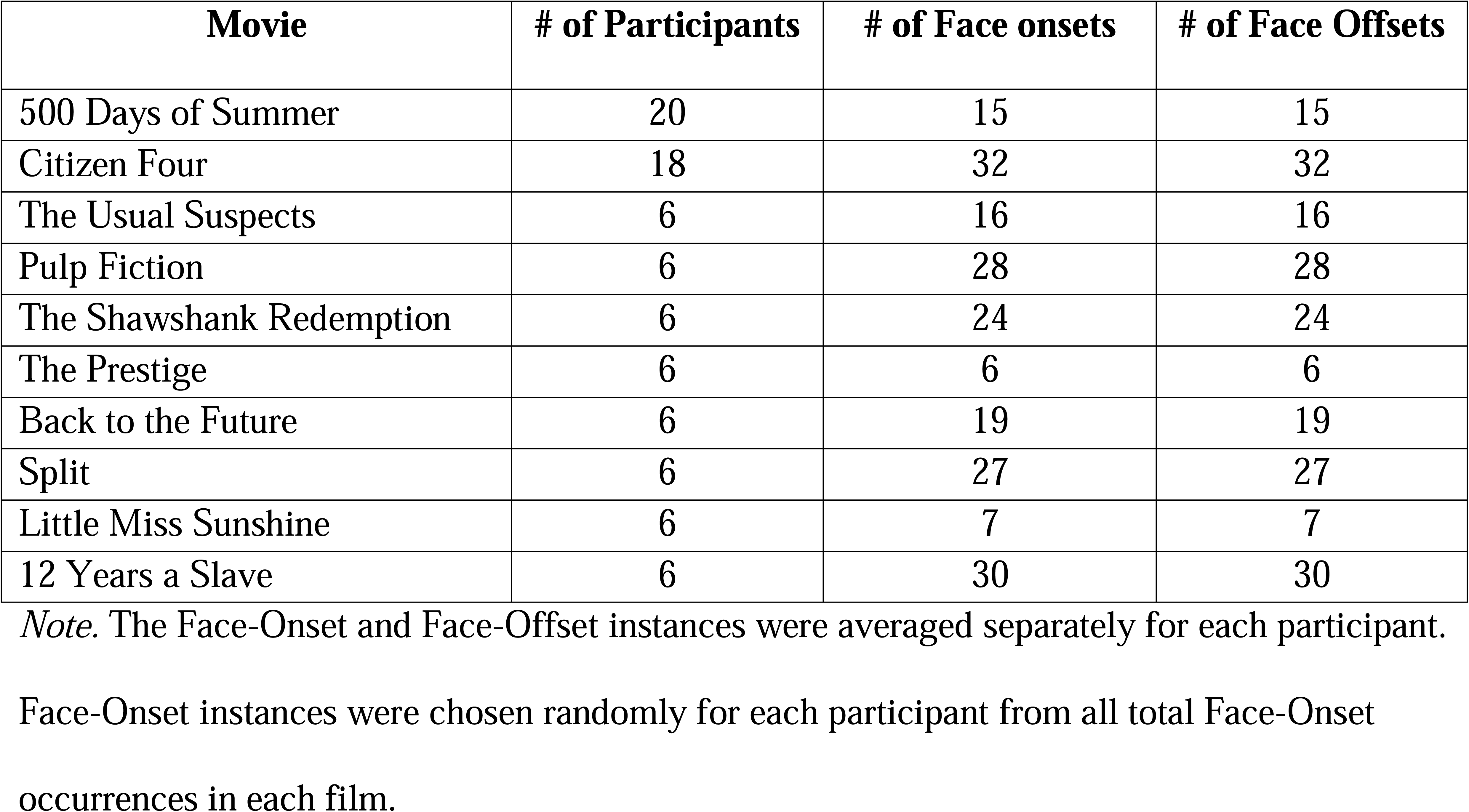
Number of Face-Onset and Face-Offset Instances Selected for Each Film

| Movie | # of Participants | # of Face onsets | # of Face Offsets |
| --- | --- | --- | --- |
| 500 Days of Summer | 20 | 15 | 15 |
| Citizen Four | 18 | 32 | 32 |
| The Usual Suspects | 6 | 16 | 16 |
| Pulp Fiction | 6 | 28 | 28 |
| The Shawshank Redemption | 6 | 24 | 24 |
| The Prestige | 6 | 6 | 6 |
| Back to the Future | 6 | 19 | 19 |
| Split | 6 | 27 | 27 |
| Little Miss Sunshine | 6 | 7 | 7 |
| 12 Years a Slave | 6 | 30 | 30 |

*Note.* The Face-Onset and Face-Offset instances were averaged separately for each participant.
Face-Onset instances were chosen randomly for each participant from all total Face-Onset occurrences in each film.

### FMRI Data Restructuring

FMRI data restructuring was performed using in-house python code (https://github.com/ekstrandlab/spatiotemporal_clustering_4d_fmri) to extract patterns of activation occurring 10-seconds after onset, and 10-seconds after offset, of human faces in audiovisual movies. To isolate fMRI activation associated with our events of interest and account for the hemodynamic lag, we extracted fMRI data acquired four seconds after stimulus onset (similar to Wittkuhn & Schuck, 2021). The processing pipeline was as follows: for each Face-onset and Face-offset index we added four TRs to that index to account for the hemodynamic lag, and selected the 10 functional volumes immediately following this new index. We then averaged all the raw signal volumes for the Face-Onset instances and Face-Offset instances together to obtain an estimate of the response amplitude for each time point separately for each participant. This resulted in two final 10 × 64 × 76 × 64 arrays for each participant reflecting the average response after 10 seconds to randomly chosen Face-Onset instances and the Face-Offset instances. We then created a new 4D file for both Face-onset and Face-offset conditions separately by stacking the averaged functional files for all of the participants, resulting in an 86 (number of participants) × 10 (volumes, TR = 1s) × 64 (voxels in x-dimension) × 76 (voxels in y dimension) × 64 (voxels in z dimension) matrix.

### Data Analysis

To identify significant spatiotemporal clusters of activation from the averaged raw HRF images, we used in-house python code, available on Github (https://github.com/ekstrandlab/spatiotemporal_clustering_4d_fmri). Specifically, to examine significant differences between Face-onset and Face-offset, we used the *spatio_temporal_cluster_1samp_test* function (Maris & Oostenveld, 2007; Sassenhagen & Draschkow, 2019) from the MNE-python stats module (Gramfort, 2013). We subtracted the Face-Offset matrix from the Face-Onset matrix to examine the difference between paired samples between the two conditions. To define adjacency between the spatial vertices in the data, we used the *combine_adjacency* function from the MNE-python stats module to combine regular lattice adjacency matrices for the x, y, and z spatial dimensions. For the time dimension, only time points directly next to each other were considered adjacent. Significant spatiotemporal clusters were corrected for multiple comparisons using permutations (number of permutations = 5000) and a cluster forming threshold corresponding to a *p*-value of 0.001. Significant clusters were corrected for multiple comparisons using a significance threshold of *p* < .05. The resulting *t-*statistic images were then masked with the significant clusters identified using permutation testing. Results were transformed into surface space for visualization purposes only.

## Results

### Face-onset vs. Face-offset paired samples *t*-test

#### Face-onset > Face-offset

Results from the Face-onset > Face-offset condition identified 16 significant clusters of activation across the 10 timepoints (see Figures 1 and 2 and Supplementary Movie 1; warm colours). This included five clusters in the bilateral occipital cortices, one cluster extending across the left lateral occipital cortex, four clusters in the bilateral fusiform gyri, five clusters in the bilateral superior temporal sulci, and one cluster extending from the right fusiform gyrus into the right superior temporal gyrus. Significant activation in the right FFA began at 2 seconds (s) and persisted until 10s. Significant activation of the left fusiform gyrus was found at 3s, persisting until 9s. Activation of the right OFA was found at 2s and continued until 8s, while activation of the left OFA began at 2s and continued until 7s. Significant activation of the bilateral LOC was found from 3s to 10s. Activation of the bilateral pSTS started at 3s and continued until 10s. Activation of the aSTS started as early as 4s and persisted until 10s. Activation of the occipital cortices was found from 3s to 10s.

**Figure 1.**
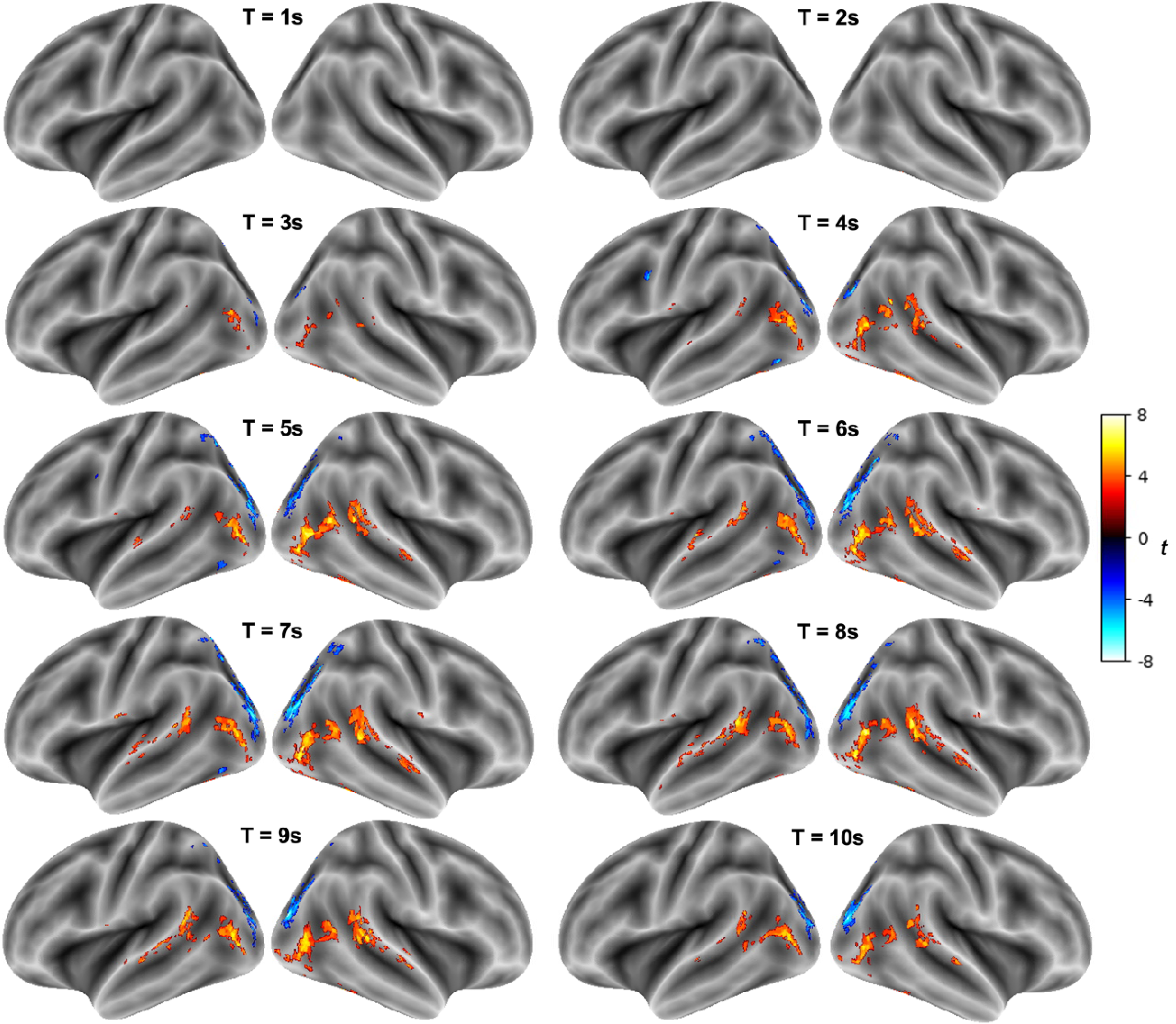
Significant activation for the Face-onset vs. Face-offset contrast over 10 seconds on the lateral surface of the brain *Note.* Significant activation for the Face-onset > Face-offset contrast are shown in warm colors and Face-offset > Face-onset is shown in cool colors. Significant clusters were corrected for multiple comparisons using permutation testing (number of permutations = 5000) and a cluster forming threshold of *p* < 0.001. Cluster significance was assessed at *p* < 0.05. Time (T) is in seconds (s). The left hemisphere is shown on the left.

**Figure 2.**
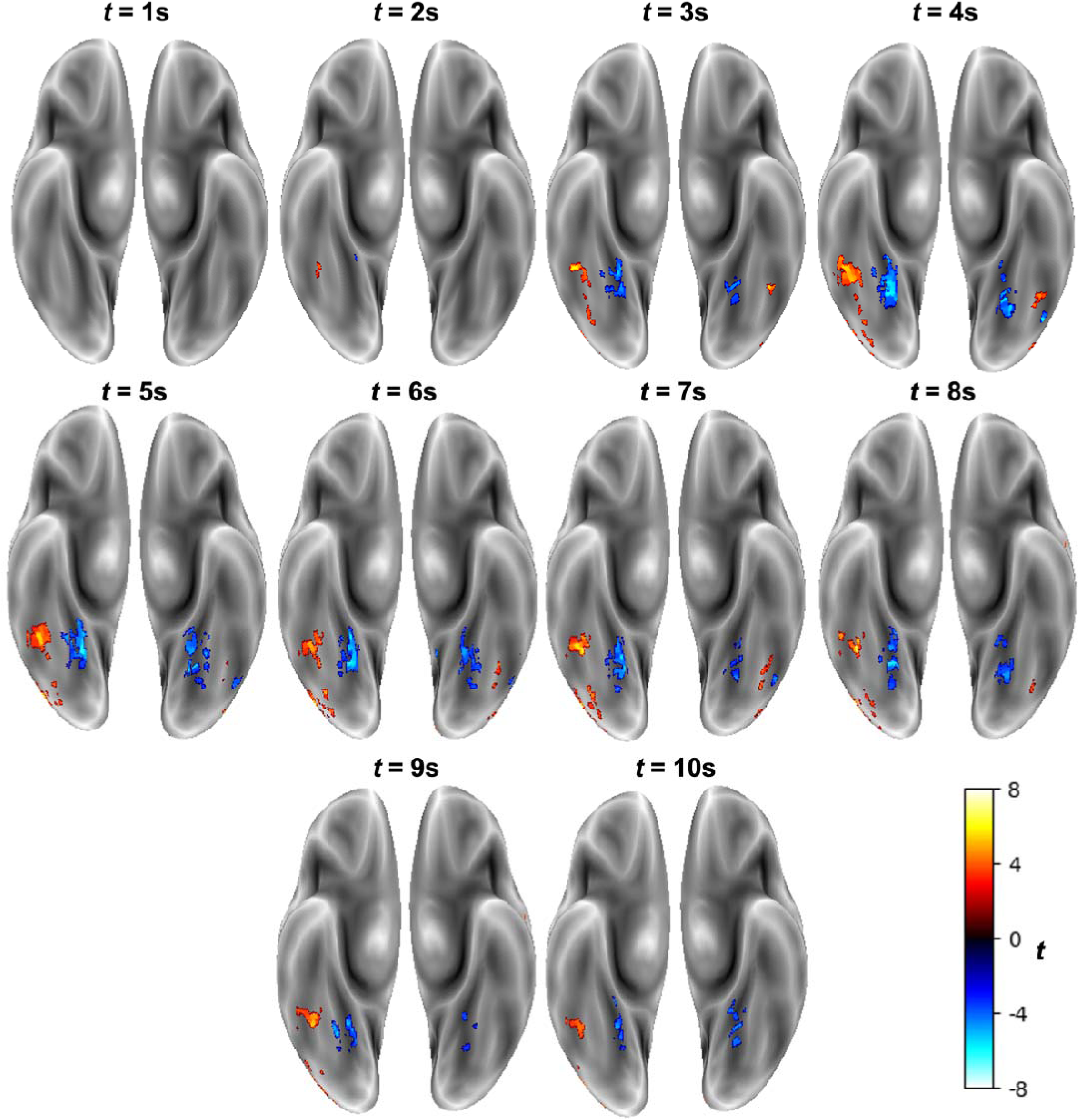
Significant activation for the Face-onset vs. Face-offset contrast over 10 seconds on the inferior surface of the brain *Note.* Significant activation for the Face-onset > Face-offset contrast are shown in warm colors and Face-offset > Face-onset is shown in cool colors. Significant clusters were corrected for multiple comparisons using permutation testing (number of permutations = 5000) and a cluster forming threshold of *p* < 0.001. Cluster significance was assessed at *p* < 0.05. Time (T) is in seconds (s). The left hemisphere is shown on the right.

#### Face-offset > Face-onset

Results from the Face-offset > Face-onset contrast identified 14 significant clusters of activation across the 10 timepoints (see Figures 1 and 2 and Supplementary Movie 1; cool colours). This included four clusters in the bilateral fusiform gyri, one cluster in the left lingual gyrus, one cluster at the border of the left precuneus, intracalcarine cortex, and lingual gyrus, six clusters in the LOCs (three superior, two extending from inferior to superior, and one inferior), one cluster in the right superior LOC/superior parietal lobule, and one cluster in the left precentral gyrus. Activation of the right lingual gyrus was found at 2s, extending into the medial fusiform cortex until 10s. Activation of the left medial fusiform cortex was found from 3s to 10s. Activation of the left precuneus/intracalcarine cortex/lingual gyrus was found at 6s. Activation of the bilateral LOCs was found from 3s to 10s, extending dorsally over time until 9s. Activation in the left precentral gyrus was found from 4s to 5s.

### Temporal dynamics associated with Face-onset

To examine the average BOLD time course associated with each cluster for the Face-onset > Face-offset contrast, we created a 3D mask of the full extent of each cluster and used this mask to characterize the average Face-onset BOLD activation for all participants in that cluster for each of our 10 timepoints. Figures 3, 4, and 5 show the full extent of our 4D clusters in 3D space with the average BOLD amplitude in that cluster at each timepoint inset for each cluster on inferior, lateral, and medial surfaces of the brain (respectively).

**Figure 3.**
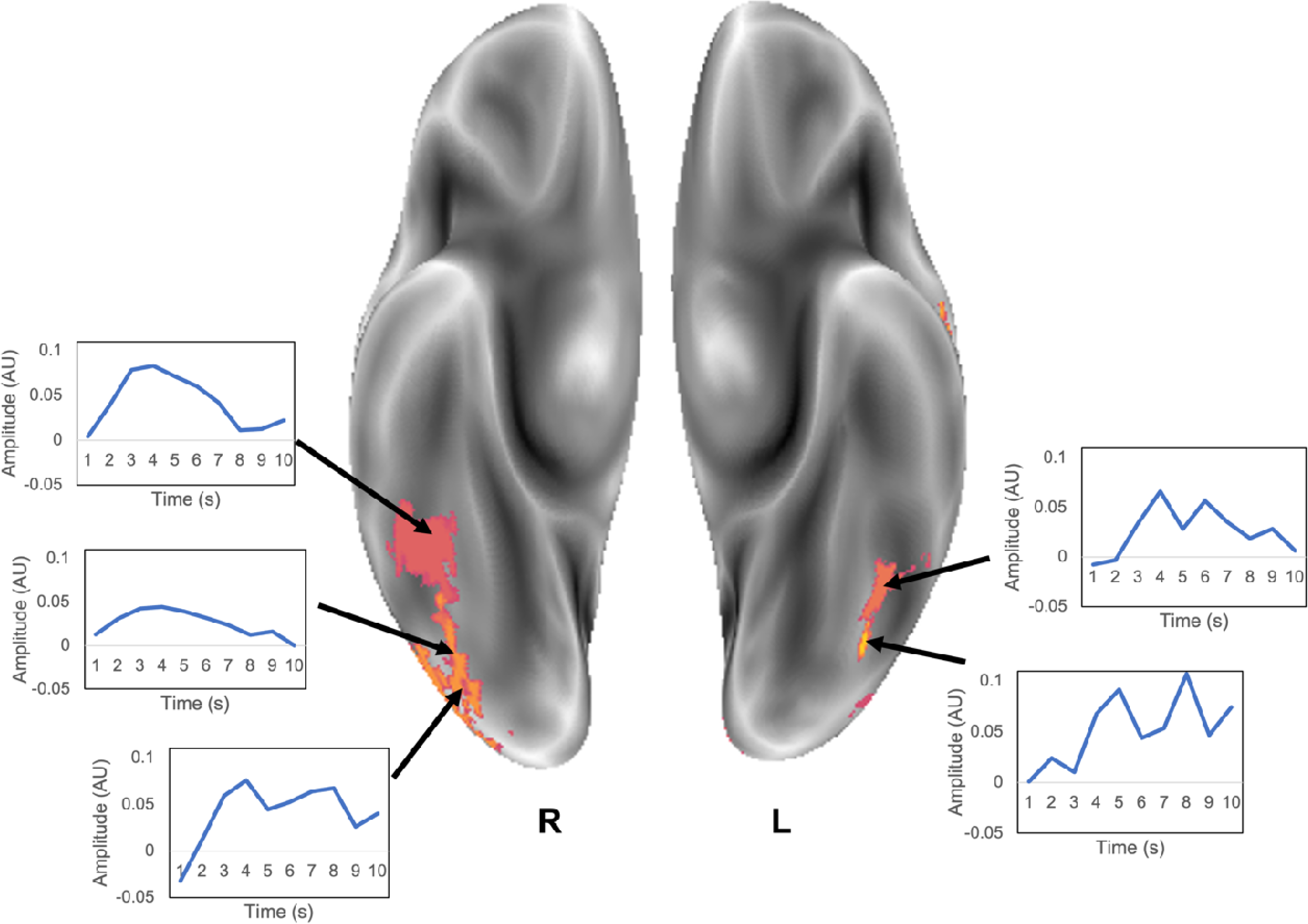
clusters across all timepoints for the Face-onset > Face-offset contrast on the inferior surface of the brain. Average response amplitude plots for each time point are inset for each cluster. *Note.* AU = Arbitrary units. Time is in seconds.

**Figure 4.**
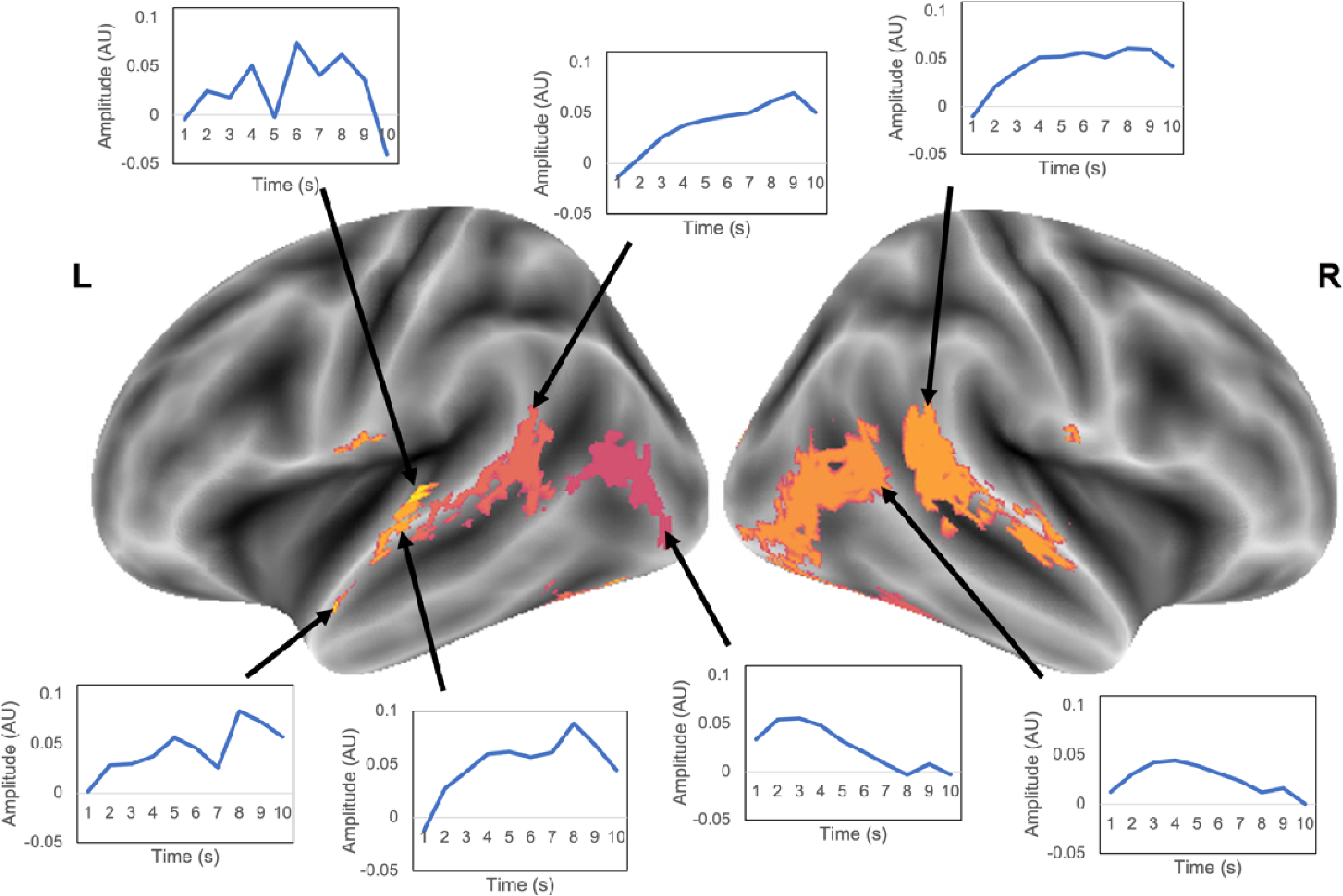
Significant clusters across all timepoints for the Face-onset > Face-offset contrast on the lateral surface of the brain. Average response amplitude plots for each time point are inset for each cluster. *Note.* AU = Arbitrary units. Time is in seconds.

**Figure 5.**
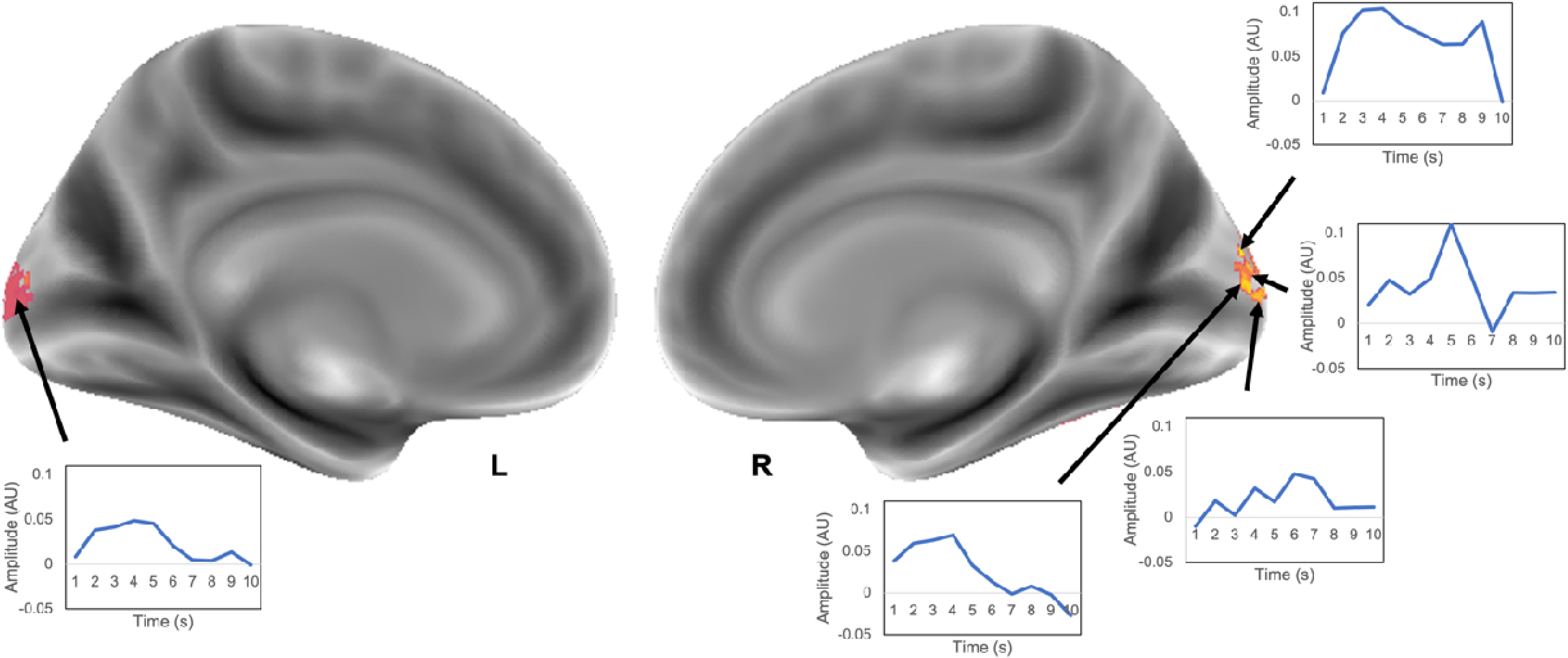
Significant clusters across all timepoints for the Face-onset > Face-offset contrast on the medial surface of the brain. Average response amplitude plots for each time point are inset for each cluster. *Note.* AU = Arbitrary units. Time is in seconds.

## Discussion

Overall, results from this study showed that the 4D spatiotemporal clustering method proposed here can extract clusters of fMRI activation over time that closely resemble the known spatiotemporal pattern of human face processing without the need to model a hemodynamic response function. Results from this study showed that onset of faces in naturalistic stimuli is associated with early activation of the right FFA, followed by activation of the bilateral occipital poles, left FFA, bilateral OFA, bilateral LOC, and right pSTS. Further, we showed that activation along the STS spread from posterior to anterior over time. Overall, the 4D clusters of activation related to face onset in audiovisual movies showed the characteristic posterior to anterior spread of activation along the ventral stream hypothesized based on previous literature (e.g., Fan et al., 2020). Examination of the average BOLD response following face onset at each timepoint for the full extent of each 4D cluster projected into 3D space showed variable patterns of activation that did not closely follow the canonical hemodynamic response profile. Together, these results provide proof of concept of the utility of this technique to provide exciting insight into temporal dynamics of BOLD activation associated with specific events in naturalistic stimuli that is not constrained by specific models of the hemodynamic response function.

Results from the Face-offset > Face-onset contrast showed a spread of activation along the dorsal stream, starting in the inferior LOCs and extending to the superior parietal lobules over time. We also found activation of the medial fusiform cortex that persisted over the duration of our timepoints in the parahippocampal place area, which is integral in scene perception. While we did not control for the type of objects present in the Face-offset condition (other than that a face was not present), it is likely that a dominant category of stimuli that is present in the face-offset instances are scenes/places. Thus, this research also indirectly supports other research using naturalistic stimuli that show activation of the parahippocampal place area in naturalistic stimuli during scene processing (Häusler et al., 2022). Future research should seek to disentangle the object categories present in the Face-offset condition to further characterize the temporal dynamics associated with different stimulus categories.

A major advantage of this approach is its relative simplicity and interpretability compared to other attempts at modeling events in naturalistic stimuli, particularly when the events of interest are known to the researcher. Previous research on fMRI activation related to events in audiovisual movies has provided insight into static activation maps associated with specific object categories (e.g., scenes/places; Häusler et al., 2022). However, with our approach, it is possible to create time-varying visualizations of the BOLD signal in the brain to draw inferences about underlying activity patterns (in this case, how face processing unfolds over time).

Importantly, our technique uses the complex nature of naturalistic stimuli to its advantage by allowing unrelated factors to be balanced across a large number of variable events in the movie. Further, previous attempts at modeling the BOLD signal in 4D have relied on multiple task-based fMRI runs for each participant (e.g., Ledberg et al., 2001) or convolution of the hemodynamic response function (e.g., Parmar & Walden, 2022), which are not ideal for naturalistic stimuli. While more complex methods focused on unraveling the dynamical organization of the brain using fMRI have been developed (e.g., Topological Data Analysis; Saggar et al., 2018), they are difficult to map back into “brain space” for visualization and interpretation. Thus, our technique provides a simple and behaviorally relevant snapshot of dynamic brain activation associated with time locked stimulus events.

In addition to its interpretability, this technique is also highly flexible. While we show only a specific example of how this analysis can be used (i.e., to examine audiovisual face processing within participants), it can in theory be applied to any event(s) of interest. In this way, researchers can ask novel and exciting questions through carefully designed contrasts, taking advantage of the variability within naturalistic stimuli. For example, by selecting instances where an actor’s face is in the same direction vs. facing different directions, viewpoint invariance in naturalistic stimuli could be assessed. This analysis could also be applied to task-based experimental designs to provide exciting insight into temporal dynamics of fMRI in more traditional experimental designs. Researchers can also examine between group differences by changing their test statistic for spatiotemporal clustering, and can also look at higher level interactions between variables of interest using ANOVA tests. Further, this research provides a jumping off point for higher level analyses focused on feature extraction by providing information about characteristic activation patterns associated with specific events, that can then be modeled in more complex analyses.

The ability to examine brain activation in 4D may offer exciting insight into group differences in neural activation that cannot be examined using traditional analyses. Previous research has suggested that many disorders, such as schizophrenia (Damaraju et al., 2014), bipolar disorder (Rashid et al., 2014), and dementia (Sourty et al., 2016), are associated with aberrant neural dynamics that are difficult to characterize with static analytic techniques. Indeed, 4D deep learning models have shown important promise for detecting pathology (e.g., Alzheimer’s disease; Kazemi & Houghten, 2018; Li et al., 2020), outperforming models based on functional connectivity, 2D, and 3D fMRI data. However, a limitation of deep learning models is that it is difficult to extract and interpret information about what features of the data are relevant for classification. Thus, the 4D spatiotemporal clustering technique described here may provide a valuable tool for characterizing and localizing aberrant neural dynamics in clinical populations.

One limitation of the current technique is that the temporal dimension is reliant on the sluggish BOLD signal and includes clustering over time, and thus does not reflect the actual speed of processing. Therefore, the focus of this analysis should be on examining the overall pattern of activation. Another limitation of the stimuli used in this study is the discrepancy between the number of Face-offset and Face-onset occurrence in each of the films. Because these were feature length movies, there were much fewer instances where a face was not on the screen for more than 10 seconds, and these scenes were often of shorter duration than the scenes where faces were on the screen. Therefore, while we were able to randomly choose face instances to average, we were unable to do this for the Face-offset instances, which may result in increased activation specific to the scenes, which would be the similar for each participant who watched the same movie. However, by using multiple movies in our analysis the effects shown here cannot be attributed solely to scene-specific activation that is consistent across participants.

Overall, this novel 4D spatiotemporal clustering technique can provide exciting insight into the dynamic functional architecture of the brain in naturalistic settings using fMRI. In this work, we were able to extract 4D clusters associated with face processing events in audiovisual movies that closely match the known pattern of spatiotemporal dynamics of human face processing. Beyond face processing, this technique can provide a window into how functional processes unfold over time and space by distilling complex neuroimaging data into a simple and behaviorally relevant representation that has a myriad of implications for researchers and clinicians alike. In conclusion, the 4D spatiotemporal clustering technique for fMRI data presented here has the potential to provide a new and exciting window into dynamic functional processing in both health and disease. Further, it may offer valuable insight into dynamic functional differences associated with pathology not readily apparent in static analyses. We hope that the current work will serve as an impetus for future research focused on uncovering the dynamic functional architecture of the human brain using fMRI.

## References

Aliko, S., Jiawen Huang, Gheorghiu, F., Meliss, S., & Skipper, J. I. (2021). Naturalistic Neuroimaging Database [Data set]. Openneuro. https://doi.org/10.18112/OPENNEURO.DS002837.V2.0.0

Bai, B., Kantor, P., & Shokoufandeh, A. (2007). Effectiveness of the Finite Impulse Response Model in Content-Based fMRI Image Retrieval. In N. Ayache, S. Ourselin, & A. Maeder (Eds.), Medical Image Computing and Computer-Assisted Intervention – MICCAI 2007 (Vol. 4792, pp. 742–750). Springer Berlin Heidelberg. https://doi.org/10.1007/978-3-540-75759-7_90

Ben-Yakov, A., Honey, C. J., Lerner, Y., & Hasson, U. (2012). Loss of reliable temporal structure in event-related averaging of naturalistic stimuli. NeuroImage, 63(1), 501–506. https://doi.org/10.1016/j.neuroimage.2012.07.008

Collins, J. A., & Olson, I. R. (2014). Beyond the FFA: The role of the ventral anterior temporal lobes in face processing. Neuropsychologia, 61, 65–79. https://doi.org/10.1016/j.neuropsychologia.2014.06.005

Cox, R. W. (1996). AFNI: Software for Analysis and Visualization of Functional Magnetic Resonance Neuroimages. Computers and Biomedical Research, 29(3), 162–173. https://doi.org/10.1006/cbmr.1996.0014

Damaraju, E., Allen, E. A., Belger, A., Ford, J. M., McEwen, S., Mathalon, D. H., Mueller, B. A., Pearlson, G. D., Potkin, S. G., Preda, A., Turner, J. A., Vaidya, J. G., Van Erp, T. G., & Calhoun, V. D. (2014). Dynamic functional connectivity analysis reveals transient states of dysconnectivity in schizophrenia. NeuroImage: Clinical, 5, 298–308. https://doi.org/10.1016/j.nicl.2014.07.003

Dawel, A., Miller, E. J., Horsburgh, A., & Ford, P. (2021). A systematic survey of face stimuli used in psychological research 2000–2020. Behavior Research Methods, 54(4), 1889– 1901. https://doi.org/10.3758/s13428-021-01705-3

Duchaine, B., & Yovel, G. (2015). A Revised Neural Framework for Face Processing. Annual Review of Vision Science, 1(1), 393–416. https://doi.org/10.1146/annurev-vision-082114-035518

Fan, X., Wang, F., Shao, H., Zhang, P., & He, S. (2020). The bottom-up and top-down processing of faces in the human occipitotemporal cortex. ELife, 9, e48764. https://doi.org/10.7554/eLife.48764

Freiwald, W., Duchaine, B., & Yovel, G. (2016). Face Processing Systems: From Neurons to Real-World Social Perception. Annual Review of Neuroscience, 39(1), 325–346. https://doi.org/10.1146/annurev-neuro-070815-013934

Gollo, L. L., Zalesky, A., Hutchison, R. M., van den Heuvel, M., & Breakspear, M. (2015). Dwelling quietly in the rich club: Brain network determinants of slow cortical fluctuations. Philosophical Transactions of the Royal Society B: Biological Sciences, 370(1668), 20140165. https://doi.org/10.1098/rstb.2014.0165

Goutte, C., Nielsen, F. A., & Hansen, K. H. (2000). Modeling the hemodynamic response in fMRI using smooth FIR filters. IEEE Transactions on Medical Imaging, 19(12), 1188– 1201. https://doi.org/10.1109/42.897811

Gramfort, A. (2013). MEG and EEG data analysis with MNE-Python. Frontiers in Neuroscience, 7. https://doi.org/10.3389/fnins.2013.00267

Häusler, C. O., Eickhoff, S. B., & Hanke, M. (2022). Processing of visual and non-visual naturalistic spatial information in the “parahippocampal place area.” Scientific Data, 9(1), 147. https://doi.org/10.1038/s41597-022-01250-4

Kazemi, Y., & Houghten, S. (2018). A deep learning pipeline to classify different stages of Alzheimer’s disease from fMRI data. 2018 IEEE Conference on Computational Intelligence in Bioinformatics and Computational Biology (CIBCB), 1–8. https://doi.org/10.1109/CIBCB.2018.8404980

Ledberg, A., Fransson, P., Larsson, J., & Petersson, K. M. (2001). A 4D approach to the analysis of functional brain images: Application to FMRI data. Human Brain Mapping, 13(4), 185–198. https://doi.org/10.1002/hbm.1032

Li, W., Lin, X., & Chen, X. (2020). Detecting Alzheimer’s disease Based on 4D fMRI: An exploration under deep learning framework. Neurocomputing, 388, 280–287. https://doi.org/10.1016/j.neucom.2020.01.053

Maris, E., & Oostenveld, R. (2007). Nonparametric statistical testing of EEG- and MEG-data. Journal of Neuroscience Methods, 164(1), 177–190. https://doi.org/10.1016/j.jneumeth.2007.03.024

Parmar, H., & Walden, E. (2022). Visualization of the Dynamic Brain Activation Pattern during a Decision-Making Task. Brain Sciences, 12(11), 1468. https://doi.org/10.3390/brainsci12111468

Pitcher, D., & Ungerleider, L. G. (2021). Evidence for a Third Visual Pathway Specialized for Social Perception. Trends in Cognitive Sciences, 25(2), 100–110. https://doi.org/10.1016/j.tics.2020.11.006

Polimeni, J. R., & Lewis, L. D. (2021). Imaging faster neural dynamics with fast fMRI: A need for updated models of the hemodynamic response. Progress in Neurobiology, 207, 102174. https://doi.org/10.1016/j.pneurobio.2021.102174

Rashid, B., Damaraju, E., Pearlson, G. D., & Calhoun, V. D. (2014). Dynamic connectivity states estimated from resting fMRI Identify differences among Schizophrenia, bipolar disorder, and healthy control subjects. Frontiers in Human Neuroscience, 8. https://doi.org/10.3389/fnhum.2014.00897

Saggar, M., Sporns, O., Gonzalez-Castillo, J., Bandettini, P. A., Carlsson, G., Glover, G., & Reiss, A. L. (2018). Towards a new approach to reveal dynamical organization of the brain using topological data analysis. Nature Communications, 9(1), 1399. https://doi.org/10.1038/s41467-018-03664-4

Sassenhagen, J., & Draschkow, D. (2019). Cluster_Jbased permutation tests of MEG/EEG data do not establish significance of effect latency or location. Psychophysiology, 56(6), e13335. https://doi.org/10.1111/psyp.13335

Schultz, J., & Pilz, K. S. (2009). Natural facial motion enhances cortical responses to faces. Experimental Brain Research, 194(1), 465–475. https://doi.org/10.1007/s00221-009-1721-9

Sonkusare, S., Breakspear, M., & Guo, C. (2019). Naturalistic Stimuli in Neuroscience: Critically Acclaimed. Trends in Cognitive Sciences, 23(8), 699–714. https://doi.org/10.1016/j.tics.2019.05.004

Sourty, M., Thoraval, L., Roquet, D., Armspach, J.-P., Foucher, J., & Blanc, F. (2016). Identifying Dynamic Functional Connectivity Changes in Dementia with Lewy Bodies

Based on Product Hidden Markov Models. Frontiers in Computational Neuroscience, 10. https://doi.org/10.3389/fncom.2016.00060

Taylor, P. A., Chen, G., Glen, D. R., Rajendra, J. K., Reynolds, R. C., & Cox, R. W. (2018). *FMRI processing with AFNI: Some comments and corrections on “Exploring the Impact of Analysis Software on Task fMRI Results”* [Preprint]. Neuroscience. https://doi.org/10.1101/308643

Wittkuhn, L., & Schuck, N. W. (2021). Dynamics of fMRI patterns reflect sub-second activation sequences and reveal replay in human visual cortex. Nature Communications, 12(1), 1795. https://doi.org/10.1038/s41467-021-21970-2

